# Subclinical infection of macaques and baboons with a baboon simartevirus

**DOI:** 10.1101/397430

**Authors:** Connor Buechler, Matthew Semler, David A. Baker, Christina Newman, Joseph P. Cornish, Deborah Chavez, Bernadette Guerra, Robert Lanford, Kathy Brasky, Jens H. Kuhn, Reed F. Johnson, David H. O’Connor, Adam L. Bailey

## Abstract

Simarteviruses (*Arteriviridae*: *Simartevirus*) are commonly found at high titers in the blood of African monkeys but do not cause overt disease in these hosts. In contrast, simarteviruses cause severe disease in Asian macaques upon accidental or experimental transmission. Here, we sought to better understand the host-dependent drivers of simartevirus pathogenesis by infecting olive baboons (n=4) and rhesus macaques (n=4) with the simartevirus Southwest baboon virus 1 (SWBV-1). Surprisingly, none of the animals in our study showed signs of disease following SWBV-1 inoculation. Three animals (two rhesus monkeys and one olive baboon) became infected and sustained high levels of SWBV-1 viremia for the duration of the study. The course of SWBV-1 infection was highly predictable: plasma viremia peaked between 1×10^7^ and 1×10^8^ vRNA copies/ml at 3–10 days post-inoculation, which was followed by a relative nadir and then establishment of a stable set-point between 1×10^6^ and 1×10^7^ vRNA copies/ml for the remainder of the study (56 days). We characterized cellular and antibody responses to SWBV-1 infection in these animals, demonstrating that macaques and baboons mount similar responses to SWBV-1 infection, yet these responses are ineffective at clearing SWBV-1 infection. SWBV-1 sequencing revealed the accumulation of non-synonymous mutations in a region of the genome that corresponds to an immunodominant epitope in the simartevirus major envelope glycoprotein GP5, which likely contribute to viral persistence by enabling escape from host antibodies.

**One Sentence Summary:** Simartevirus infection has multiple disease manifestations following cross-species transmission.

**Accessible Summary/Importance:** Simarteviruses are known to infect African monkeys, such as olive baboons, without causing overt disease. In contrast, accidental infection of Asian monkeys, such as rhesus monkeys, has resulted in severe and often fatal disease. We used a simartevirus found circulating among captive olive baboons (Southwest baboon virus 1; SWBV-1) to experimentally infect both olive baboons and rhesus monkeys to model infection with the same virus in both natural and non-natural hosts. Surprisingly, neither baboons nor macaques displayed any laboratory abnormalities or signs of disease over the course of infection, despite robust SWBV-1 replication. In the accompanying study by Cornish et al., a similar experimental approach was undertaken: African patas monkeys and rhesus monkeys were infected with the simartevirus simian hemorrhagic fever virus (SHFV). In contrast to our study, SHFV caused disease in both of these hosts, albeit with much more severe disease developing in the macaques. Interestingly, we observed similar levels of immune cell activation in simartevirus-infected animals across both studies, suggesting that finer nuances of the host response, and perhaps properties of each individual simartevirus, may influences pathogenicity of these viruses in primates. Taken together, our collective findings highlight the wide clinical spectrum of simartevirus infection, ranging from highly-lethal hemorrhagic disease to persistent infection without any overt signs of disease, even in non-natural primate hosts.

## Introduction

Wild non-human primates are an important reservoir of several zoonotic pathogens, and recent surveys have shown that many harbor microbes with unknown zoonotic potential (1–4). In particular, a high proportion of wild monkeys across sub-Saharan Africa harbor arteriviruses (order: *Nidovirales*; family: *Arteriviridae*) that have a unique genomic architecture (genus: *Simartevirus*) (5, 6). In naturally-infected wild African monkeys, simarteviruses cause persistent high-titer viremia and exhibit a high degree of intra- and inter-host genetic variation without eliciting any overt signs of disease—features that may make cross-species transmission to humans more likely (5).

Wild Asian macaques (*Macaca* spp.) are not known to harbor simarteviruses, but captive macaques have been infected with simarteviruses (both experimentally and unintentionally) and develop severe and frequently fatal disease characterized by hemorrhage and multi-organ failure, with mortality often exceeding 50% (7–18). Indeed, the first two simarteviruses identified, simian hemorrhagic encephalitis virus (SHEV) and simian hemorrhagic fever virus (SHFV), were first encountered in 1964 following two seemingly connected outbreaks of viral hemorrhagic fever in captive macaques at primate facilities in Sukhumi, Georgian SSR, USSR, and the National Institutes of Health in Bethesda, Maryland, USA, respectively (7). Since the discovery of these two viruses, many other outbreaks have occurred in captive Asian macaques throughout the world, with captive co-housed African monkeys often implicated as the source of infection. Recent retrospective investigations indicate that many of these outbreaks may have been caused by viruses distinct from SHEV and SHFV, suggesting that simarteviruses as a group may be universally pathogenic to Asian macaques (19). However, not all simarteviruses are universally fatal in macaques, as crab-eating macaques (*Macaca fascicularis*) infected with the simartevirus Kibale red colobus virus 1 (KRCV-1) isolated from wild red colobus (*Procolobus rufomitratus tephrosceles*) in Uganda exhibited mild signs of disease and fully recovered upon clearance of KRCV-1 viremia (18).

We recently discovered a novel simartevirus circulating in apparently healthy olive baboons (*Papio anubis*) at the Southwest National Primate Research Center (SNPRC) in San Antonio, Texas, USA, that we termed Southwest baboon virus (SWBV-1, hereafter referred to as “SWBV”) (20). Using this virus, we sought to understand differences in simartevirus infection and disease in natural (*i.e.* baboon) versus non-natural (*i.e.* macaque) hosts that are 98% genetically identical, with the broader goal of understanding host factors that facilitate or protect against the development of viral hemorrhagic fever. Concurrently, a similar study was conducted in patas monkeys (*Erythrocebus patas*) and rhesus macaques (*macaca mulatta*) using SHFV (see accompanying article by Cornish et al.).

## Results

### Prevalence of SWBV infection at SNPRC and selection of SWBV-naïve baboons

To further define the prevalence of SWBV infection in baboons at SNPRC, we screened blood collected from 30 random baboons (19 olive baboons [*Papio anubis*]; 10 olive-yellow [*Papio cynocephalus*] hybrid baboons; and 1 hamadryas-olive [Papio *hamadryas*] hybrid baboon) as well as 6 former olive baboon cage-mates of two olive baboons known to be SWBV-infected using a SWBV-specific qRT-PCR assay described previously. Of these animals, only one olive baboon (a former cage-mate), baboon 15290, had SWBV RNA detectable in its blood at a titer of 3.4×10^5^ vRNA copies/ml of plasma, for a prevalence of roughly 3% **(Fig. 1A,B)**. Unbiased deep sequencing of this specimen did not result in the identification of any other potential pathogens (20). To minimize the likelihood of prior SWBV infection, we selected four adult male baboons from the specific-pathogen free (SPF) colony at SNPRC. These animals had tested negative for a variety of common primate pathogens, were brought into the SPF colony as infants, and reared separately from the general colony. The baboons also tested negative for SWBV infection by qRT-PCR.

**Figure 1.**
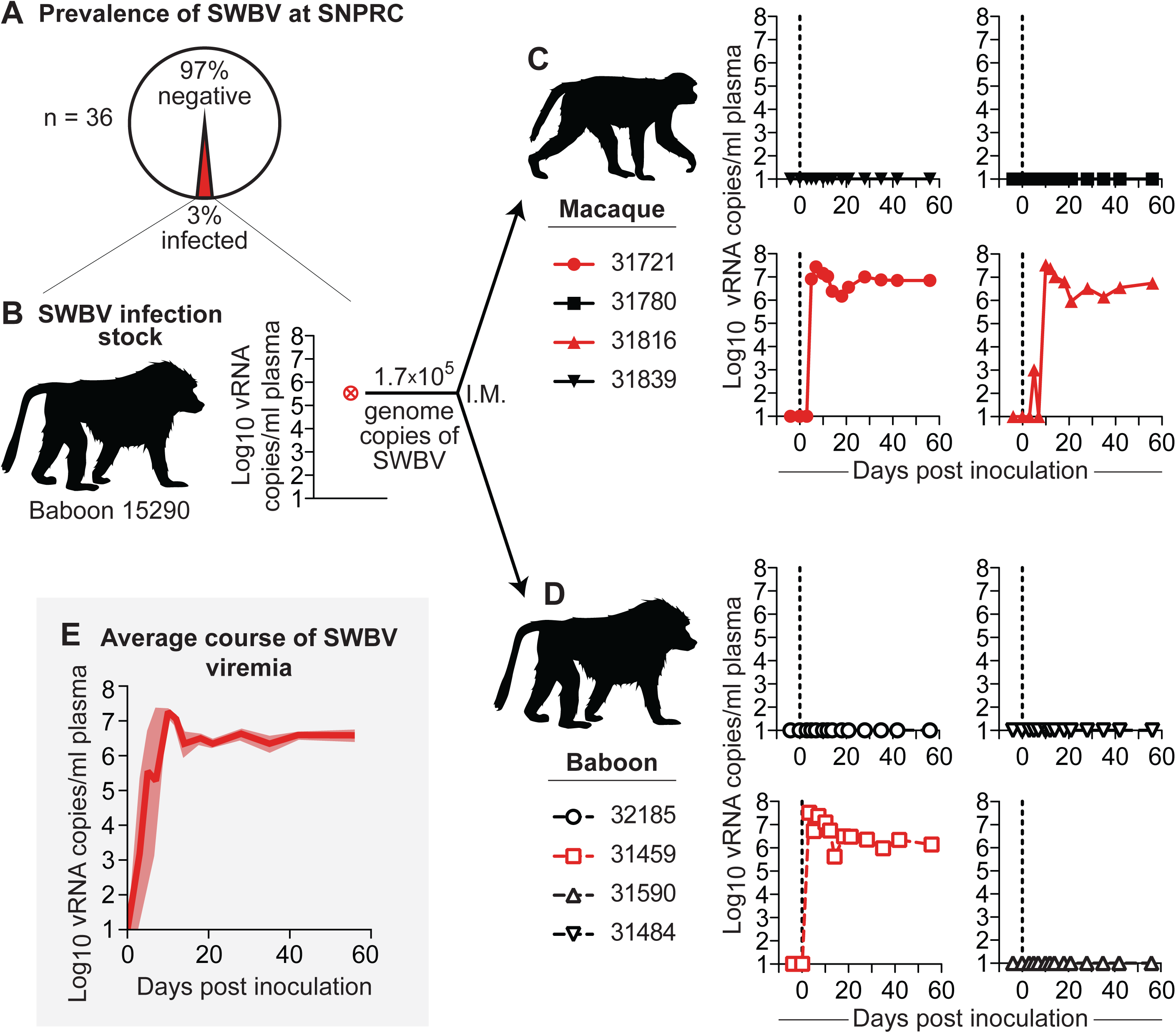
Infection of baboons and macaques with SWBV. (A) Thirty-six baboons from the specific-pathogen free colony at SNPRC were screened for SWBV infection using qRT-PCR. (B) This identified one SWBV+ baboon. Serum from this animal was used to inoculate (C) macaques (solid symbols with solid lines) and (D) baboons (open symbols with dashed lines), resulting in productive infection (red) or no infection (black) for animals from each species. Note: coloring and symbols denoting species, animal ID, and infection-status are used consistently throughout the manuscript. (E) The average course of SWBV viremia from all infected animals is in dark red, with the standard error of the mean shown in lighter red.

### The course of SWBV infection in baboons and macaques

To study differences in SWBV infection between baboons and macaques, we inoculated four olive baboons (*Papio anubis*, hereafter referred to as “baboons”) and four rhesus macaques (*Macaca mulatta*, hereafter referred to as “macaques”) intramuscularly with 500 μl of serum from baboon 15290 containing 1.7×10^5^ vRNA copies of SWBV and quantified viremia using qRT-PCR **(Fig. 1C,D)**. Five animals (two macaques and three baboon) had no detectable SWBV viremia at any time point. Three animals (two macaques and one baboon) became productively infected. Among the infected animals, the course of SWBV infection was highly predictable: plasma viremia peaked between 1×10^7^ and 1×10^8^ vRNA copies/ml at 3–10 days post-inoculation, followed by a relative nadir and then establishment of a stable set-point between 1×10^6^ and 1×10^7^ vRNA copies/ml for the remainder of the study (56 days) **(Fig. 1E)**.

### SWBV does not cause overt disease in baboons or macaques

We closely monitored all animals for signs of disease using a comprehensive clinical scoring rubric. At no time-point did any animal become clinically ill (score ≥5), nor were there significant differences between SWBV-infected and uninfected animals **(Fig. 2A)**. This result is in stark contrast to past studies of simian hemorrhagic encephalitis virus (SHEV) or simian hemorrhagic fever virus (SHFV) infection in crab-eating macaques, stump-tailed macaques, and rhesus monkeys, which resulted in extremely high morbidity and mortality **(Fig. 2B)**. No significant changes from baseline vital signs, or vital sign differences between SWBV-infected and uninfected animals of either species, were observed **(Fig 2C)**.

**Figure 2.**
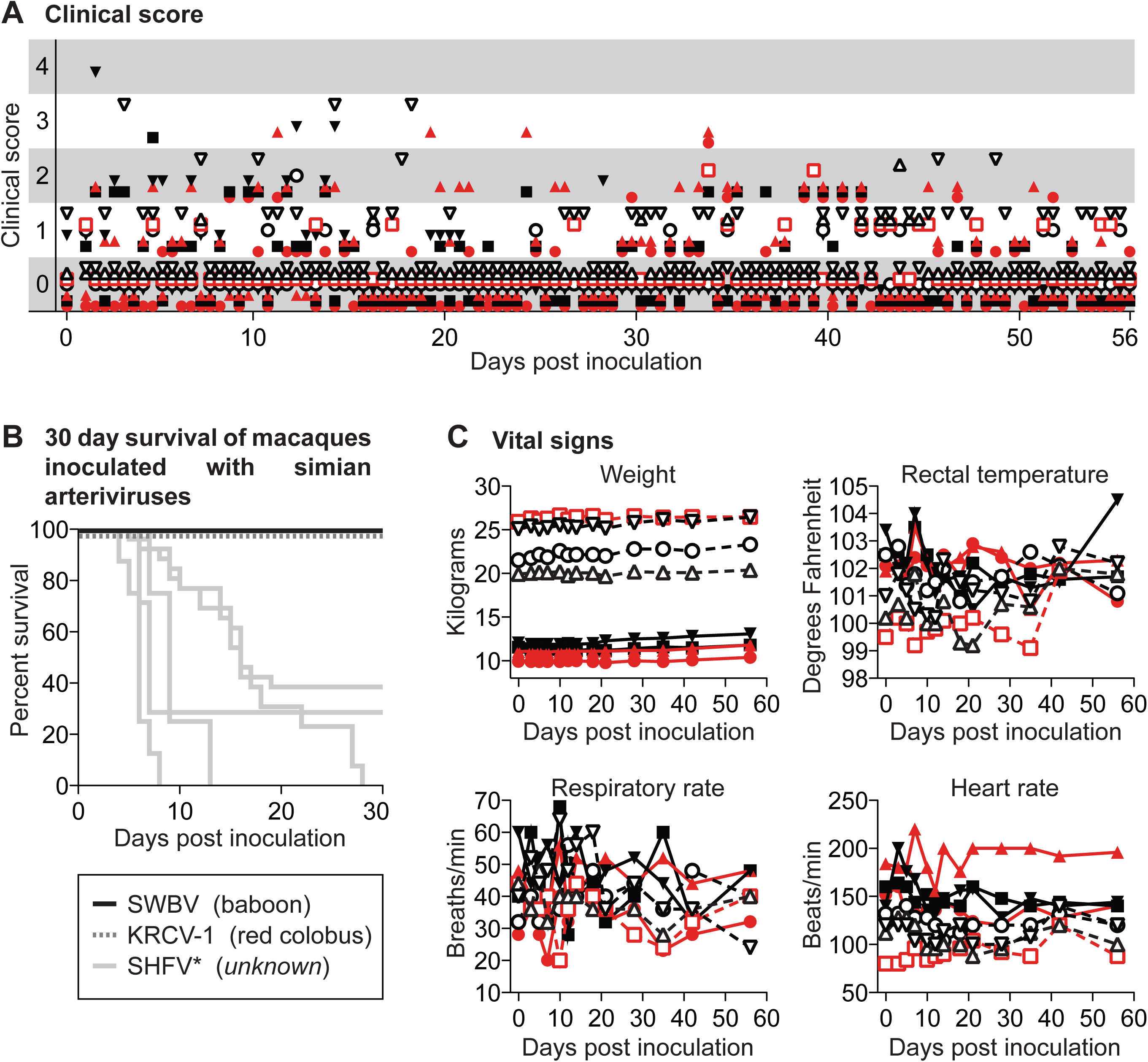
Clinical evaluation of baboons and macaques inoculated with SWBV. **(A)** Animals were evaluated for signs of disease twice per day using a comprehensive clinical scoring rubric in which a score of <5 was deemed within normal limits. **(B)** For macaques, mortality was also assessed (black line) and compared to previous studies of simartevirus infection in macaques; the virus and hosts for the simarteviruses shown are found below the graph in parentheses (SHFV has an asterisk because the natural host is not known and the sequence identity of the viruses used in some of these studies has never been confirmed). Vital signs for baboons and macaques are shown in **(C)**.

To examine more sensitive markers of infection, inflammation, and organ dysfunction, we performed a battery of laboratory tests on each animal over the course of the study, the most pertinent of which are shown **(Fig. 3)**. No significant changes in white blood cell counts, or the numbers of leukocyte subsets, were seen over the course of the experiment for either SWBV-infected or uninfected animals **(Fig. 3A)**. Additionally, in stark contrast to previous studies of SHFV in macaques, no coagulation abnormalities were observed for any of the animals, as measured by blood concentrations of fibrinogen or functional clotting assays including pro-thrombin time (PT) and partial thromboplastin time (PTT) **(Fig. 3B)**. Blood concentrations of creatinine, AST, and ALT—which serve as markers of kidney function and liver cell death, respectively—were not significantly different between SWBV-infected and uninfected animals **(Fig 3C)**.

**Figure 3.**
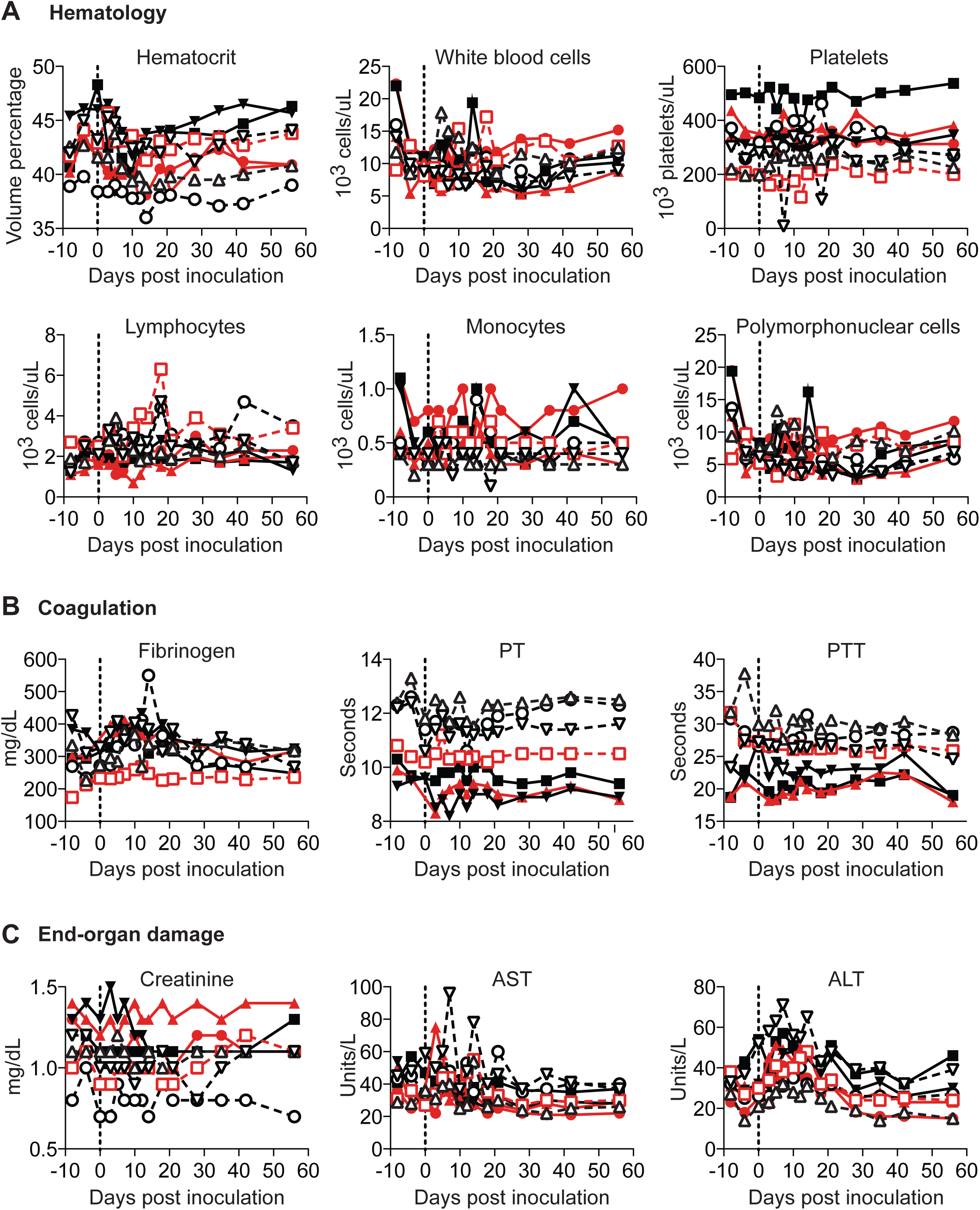
Laboratory evaluation of baboons and macaques inoculated with SWBV. Clinical chemistry, hematology, and coagulation studies were performed on blood collected from animals throughout the study.

### Arterivirus-specific adaptive immune responses in SWBV-infected macaques and baboons

To evaluate immune responses to SWBV, we assessed immune-cell activation in peripheral blood mononuclear cells (PBMC) from SWBV-inoculated baboons and macaques using flow cytometry **(Fig. 4A)**. No significant deviations from baseline levels of CD4+ T cell, CD8+ T cell, or natural killer (NK) cell activation were observed in uninfected animals (Fig. 4B). However, the percentage of activated CD4+ T cells, CD8+ T cells, and NK cells increased substantially in animals that became SWBV-viremic **(Fig. 4B)**: In Macaque 31721 and Baboon 31459, the percentage of activated immune cells peaked at day 10 post-inoculation and then quickly returned to near-baseline levels; activation of these immune-cell subsets peaked later in Macaque 31816 (between day 14 and 21) with the proportion of activated CD8+ T cells remaining elevated through day 56 post-inoculation.

**Figure 4.**
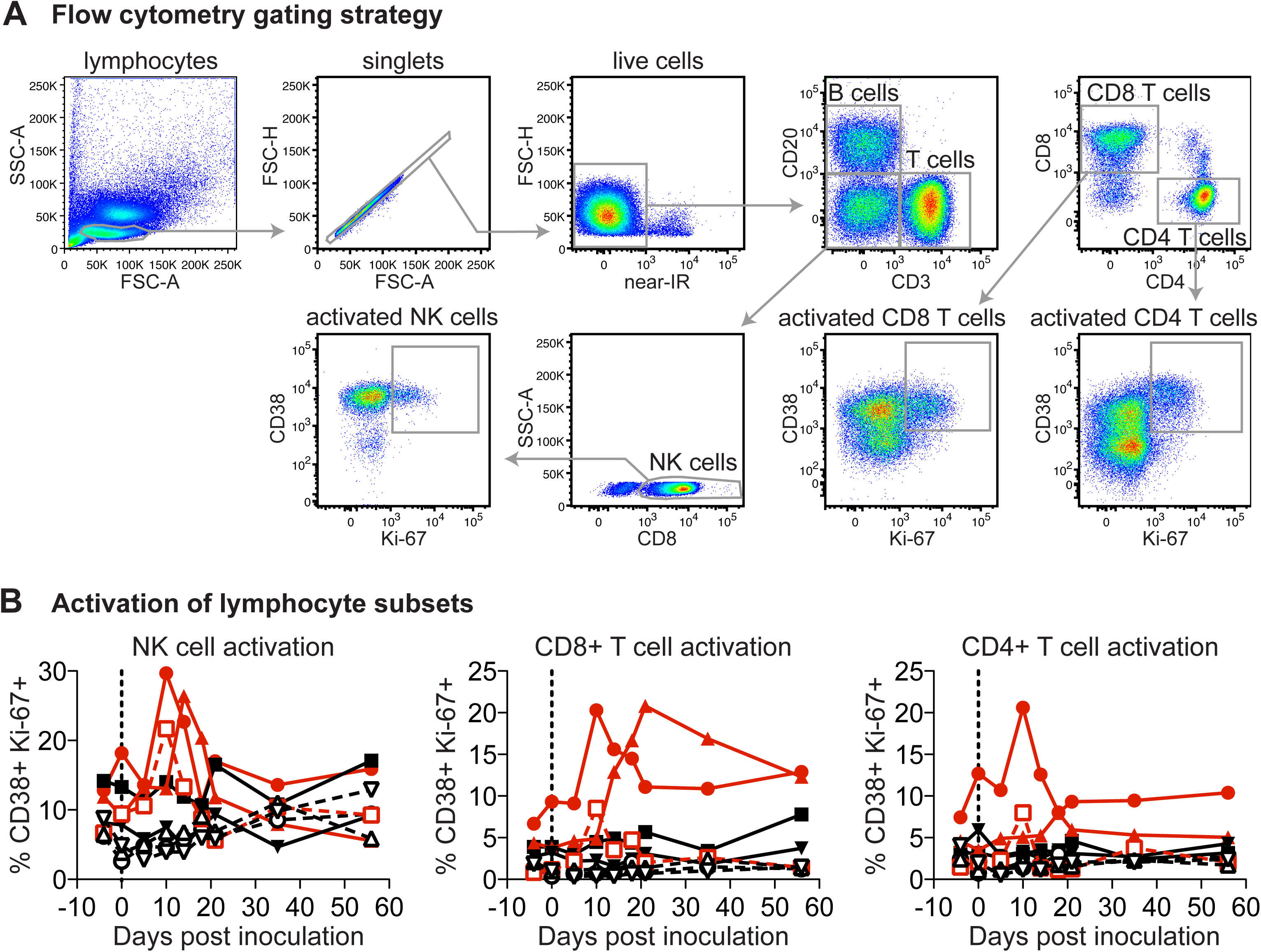
Flow cytometric analysis of peripheral blood lymphocytes from baboons and macaques inoculated with SWBV. Flow cytometry was performed on peripheral blood mononuclear cells (PBMC) that were purified and cryopreserved at the time of collection. Lymphocytes were phenotyped according to cell-surface markers and co-expression of CD38 and Ki-67 were used as a marker of cellular activation.

We also assessed antibody responses to SWBV using a microarray of 16-mer peptides that spanned the SWBV proteome with a one amino acid offset. Many arterivirus envelope glycoproteins are predicted to have relatively unstructured proteins and have been shown to elicit antibodies that recognize linear epitopes, suggesting that many of the antibodies identified by this approach would be functional (21). No SWBV-specific antibodies were detected at levels above background in any animals that were aviremic throughout the study **(Fig 5A)**. In SWBV-infected animals, SWBV-specific antibodies became detectable between 12 and 28 days post-inoculation. Although multiple envelope glycoproteins were targeted by antibodies in each of the SWBV-infected animals, common responses were focused against glycoproteins 5 and 6 (GP5 and GP6), with the most robust responses generated to GP5 **(Fig 5B).** In particular, two distinct regions/epitopes of GP5, corresponding to amino acids 28-62 and 76-111, were targeted, with the macaques mounting more robust responses to GP5_28-62_ and the baboon mounting a more robust response to GP5_76-111_.

**Figure 5.**
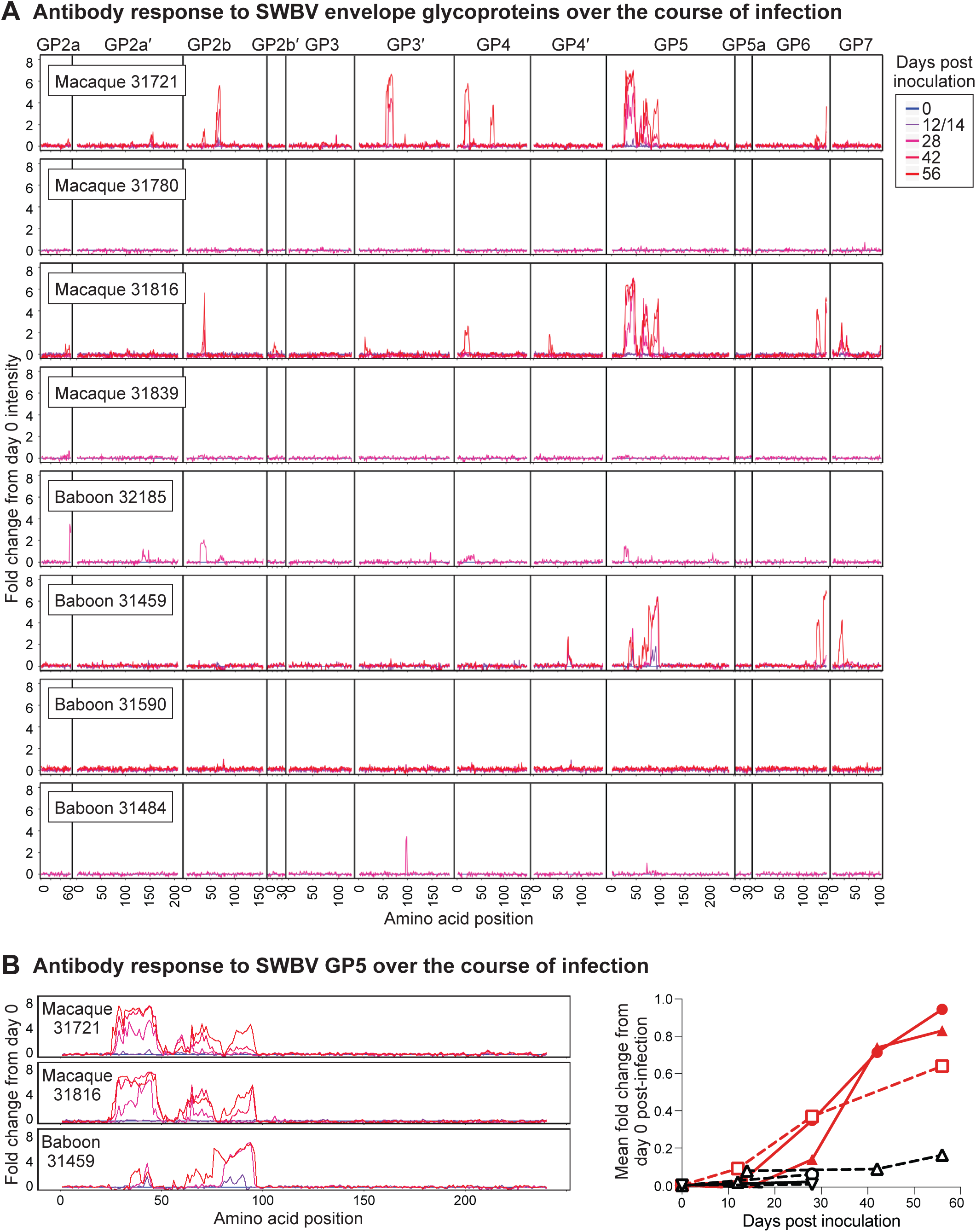
Peptide array analysis of antibodies generated by baboons and macaques inoculated with SWBV. A custom-designed array of 16-mer peptides spanning each of the SWBV proteins predicted from the SWBV inoculum nucleotide sequence was constructed and plasma from SWBV-infected animals was overlaid on this array to identify SWBV-specific antibodies. (A) Intensity of antibodies binding to peptides corresponding to the envelope glycoproteins of SWBV, normalized to the day zero post-inoculation intensity for each peptide from each animal, with day zero shown in blue and subsequent days shown in increasing shades of red. Note, numbers on the X-axis correspond to the first amino acid in the 16-mer peptide. (B) Normalized intensities over time from the GP5 protein of the three animals that became infected with SWBV; the graph to the right shows the mean fold change from day zero post-inoculation for all GP5 peptides, with SWBV-infected animals shown in red.

### Intra-host evolution of SWBV over the course of infection

We examined the genetic changes in SWBV that accumulated over the course of each animal’s infection by performing deep sequencing on plasma collected from each viremic animal. Compared to the virus used for inoculation, we found mutations in nearly every SWBV gene (Table S2), many of which were not present at detectable levels in the inoculum. However, the density of non-synonymous mutations was not evenly distributed, with many mutations occurring in the genes encoding envelope glycoproteins, particularly the GP5-encoding ORF5 gene.

## Discussion

In this study we attempted to infect baboons and macaques with a baboon arterivirus, SWBV, in order to more fully characterize the pathogenesis of simartevirus infection in natural versus non-natural hosts. Given the history of simarteviruses causing severe disease in macaques, we expected macaques infected with SWBV to develop significant disease. Surprisingly, however, SWBV resulted in clinically unapparent disease in macaques, despite high-titer and persistent viremia. While high-titer and persistent viremia is likely the most common course of simartevirus infection in naturally infected African monkeys (5), this is (to our knowledge) the first example of persistent simartevirus infection in macaques. The lack of disease observed in this study is in contrast to the accompanying study performed by Cornish et al., in which patas monkeys and rhesus macaques both developed laboratory aberrations and physical signs of disease following infection with SHFV. Therefore, it appears that simartevirus infections can be: [1] sub-clinical in African monkeys (6, 20, 22–25), [2] pathogenic in African monkeys (26–28) and Cornish et al. (2018), [3] sub-clinical in Asian-origin macaques (this study), and [4] pathogenic in Asian-origin macaques (7–18). The viral factors that that render some simarteviruses (*e.g.* SWBV) non-pathogenic but confer virulence to other simarteviruses (*e.g.* SHFV) remain to be defined, but clearly cannot be explained by host differences alone.

Analysis of immune responses to SWBV and SHFV provided no additional explanation for the range of disease observed in this study and the study by Cornish et al. In this study, SWBV infection persisted in all three animals that became infected despite SWBV-specific immune responses. Antibody responses to SWBV primarily were directed against the major envelope glycoprotein, GP5. More specifically, antibodies were raised against two N-terminal epitopes in GP5 corresponding to amino acids 28-62 and 76-111. Although the relative strength of these responses differed between the macaques and the baboon, with antibodies to the epitope corresponding to amino acids 28-62 dominating in the macaques and antibodies to the epitope corresponding to amino acids 76-111 dominating in the baboon, none of these responses was capable of achieving viral clearance. Antibody responses to the other arteriviruses have also been mapped to two N-terminal epitopes in GP5 which (29–32).

Intriguingly, the more N-terminal of these two epitopes in porcine reproductive and respiratory syndrome virus (PRRSV) is immunodominant but elicits a non-neutralizing response, referred to by some as a “decoy epitope.” (33). A similar mechanism of immune evasion may be employed by simarteviruses: indeed, we identified amino acid changes in these epitopes that would presumably abrogate antibody binding and allow SWBV to establish persistent infection by escaping antibody responses. Interestingly, the vast majority of non-synonymous mutations in the viruses from these animals were found in the sequence corresponding to the more C-terminal epitope (GP5 amino acids 76-111), further suggesting that antibodies to the more N-terminal epitope (GP5 amino acids 28-62) are non-neutralizing and do not exert a significant selective pressure on the virus **(Fig. 6)**. This also suggests that antibodies to the C-terminal epitope exert a stronger selection pressure on the virus, potentially implicating this epitope as target of neutralizing antibodies, similar to what has been observed in other arteriviruses.

**Figure 6.**
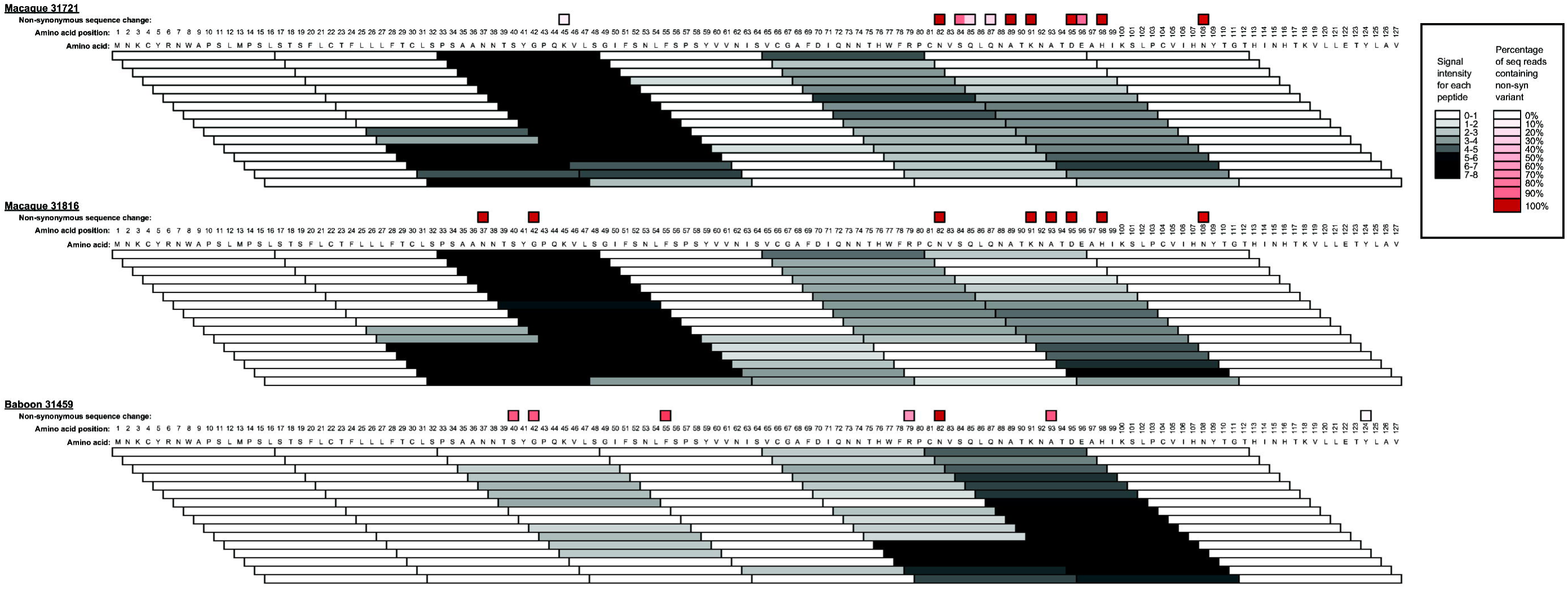
Correlation of SWBV mutations with antibody responses. Antibody binding intensity data (see Fig. 5) is shown for each peptide covering the region of interest in GP5 as a heat-mapped scale of O (white) to >7 (black). Overlayed are the non-synonymous sequencing changes corresponding to each amino acid of GP5 (see Fig. 6), with a heatmap highlighting variants contained in 20% (pink) to 100% (dark red) of deep sequencing reads. Data for each animal corresponds to analysis of the day 56 post-infection time-point with the exception of sequencing data from Baboon 31459, for which the latest timepoint (day 12 post-infection) was used.

The majority of animals in this study (5/8) did not become infected upon inoculation with SWBV. This seems unlikely to be due to pre-existing immunity from a prior SWBV infection, as this would have presumably resulted in detectable anti-SWBV antibodies and/or robust proliferation of immune cells following SWBV inoculation—neither of which were seen. Immunity due to prior or ongoing infection with a different simartevirus is also unlikely, as there does not seem to be heterologous protection between divergent simartevirus strains (18). A technical error in the inoculation process is also possible but unlikely considering that animals in each group were successfully infected. SWBV also replicated to high titers in animals that became viremic, arguing against SWBV being poorly adapted to replication in the baboon or macaque host. Therefore, it is possible that the uninfected animals were innately resistant to simartevirus infection, or that we used a dose of SWBV (1.7×10^5^ vRNA copies) that only infects 50% of animals (∼1 infectious dose 50% [ID_50_]).

Several parallels can be drawn between the simarteviruses and simian immunodeficiency viruses (SIVs): both naturally infect monkeys throughout sub-Saharan Africa and have variable propensities for causing disease upon transmission to non-natural primate hosts. Cross-species transmission of SIVs to humans and other great apes has had devastating consequences (*e.g.* the global HIV-1-pandemic), but not all SIVs have the same potential for infecting and causing disease in humans. By analogy, the consequences of simartevirus infection in humans remains unknown, and different simarteviruses may have different propensities for infecting and causing disease in humans (5). If macaques are the best proxy for assessing such variables (as they are in the case of SIV/HIV), then it is possible that simarteviruses are capable of causing a range of disease in humans including asymptomatic infection. Consequently, we believe that simartevirus research should be intensified to allow an appropriate risk assessment.

## Materials and methods

### Study design

The experimental design and protocols were agreed upon prior to initiation of the study and were not modified after the study had begun. Animal care, observations, and clinical laboratory testing were performed at the Southwest National Primate Research Center (SNPRC) in San Antonio, Texas, USA. Viral loads, flow cytometry, and data analysis were performed in batch at the Wisconsin National Primate Research Center (WNPRC), Madison, Wisconsin, USA, after conclusion of the study.

### Care and ethical use of animals

All baboons and macaques used in this study were cared for by the staff at the Southwest National Primate Research Center in accordance with the regulations and guidelines outlined in the Animal Welfare Act and the Guide for the Care and Use of Laboratory Animals. Details of this study (protocol 1539 PC, MM 0) were approved by the Texas Biomedical Research Institute’s Animal Care and Use Committee, in accordance with recommendations of the Weatherall Report.

### Clinical scoring

Animals were observed for signs of disease and scored according to the parameters described below. Scores equal to or greater than 5 required evaluation by the study veterinarian. Scores equal to or greater than 15 were deemed terminally ill. Weight loss: 0 = decrease of 0-10%; 1 = decrease of 10-20%; 2 = decrease of ≥20%. Temperature: 0 = change <2°F; 1 = change of 2-3°F; 2 = change of 3-4°F; 3= change of ≥5°F. Responsiveness: 0 = alert, responsive, normal activity, free of disease signs or exhibits only resolved/resolving disease signs; 1 = slightly diminished general activity, subdued but responds normally to external stimuli; 2 = Withdrawn, may have head down, fetal posture, hunched, reduced response to external stimuli; 8 = prostrate but able to rise if stimulated, moderate to dramatically reduced response to external stimuli; 15 = persistently prostrate, severely or completely unresponsive, may have signs of respiratory distress. Hair coat: 0 = normal appearance; 1 = rough hair coat. Respiration: 0 = normal breathing; 3 = labored; 15 = Agonal. Petechia: 0 = none; 1 = mild (1-39%); 2 = moderate (40-79%); 3 = severe (≥80%). Bleeding: 0 = none; 1 = at bleeding site; 2 = other than bleeding site. Nasal discharge: 0 = not present; 1 = present. Chow eaten: 0 = 100-25%; 1 = <25%. Food enrichment: 0 = 100-25%; 1 = <25%. Stool: 0 = normal; 1 = no stool present; 2 = diarrhea. Fluid Intake: 0 = drinking normal amounts; 1 = reduced fluid intake; 2 = no intake. Skin tenting: 0 = normal, 1 = ≥2sec.

### Virus inoculations

A SWBV stock was created for this study by aliquoting plasma collected from olive baboon 15290, which was found to be viremic with SWBV via RT-qPCR during a screen of 30 baboons at SNPRC. Macaques and baboons infected with SWBV in this study were inoculated intramuscularly with 500 μl of plasma from this animal containing 1.7 × 10^5^ vRNA copies of SWBV.

### Deep sequencing

Samples were processed for sequencing in a biosafety level 3 laboratory. For each animal, viral RNA was isolated from approximately 200 μl of plasma using the Qiagen QIAamp MinElute virus spin kit (Qiagen, Hilden, Germany), omitting carrier RNA. cDNA synthesis and PCR were performed using random hexamers (for the inoculum) or with seven primer sets (for the remaining animals, see Table S1) to amplify the entire SWBV genome in overlapping amplicons with the Superscript III One-Step RT-PCR kit with Platinum Taq Polymerase (Invitrogen, Carlsbad, CA). All amplicons from a single sample were pooled to achieve 1ng of total amplified DNA in 5 μl. Pooled amplicons (or total cDNA generated from random-hexamer amplification) were fragmented, and sequencing adaptors were added using the Nextera XT DNA Library Preparation Kit (Illumina, San Diego, CA, USA). Indexed sequencing libraries were cleaned using AMPure XP beads (Agencourt Bioscience Corporation, Beverly, MA, USA), quantified using the Qubit dsDNA HS Assay Kit with the Qubit fluorometer (Invitrogen, Carlsbad, CA, USA), and product length was measured using the Agilent 2100 Bioanalyzer High Sensitivity DNA kit (Life Technologies, Madison, WI, USA). Deep sequencing was performed on the Illumina MiSeq. Sequencing of SWBV from baboon 15290 was performed in 2016; all other sequencing was performed on the same MiSeq run in 2018. SWBV sequencing reads pertaining to this study can be found online at http://www.ncbi.nlm.nih.gov/bioproject/483240.

### Sequence read mapping and variant calling

Raw sequencing reads from baboon 15290 (the inoculum) were mapped to the previously-published SWBV-1 sequence (GenBank #KM110947) to generate an inoculum consensus sequence (GenBank #MH686150). Annotations were then transferred from KM110947 to MH686150 and checked manually. For baboon 31459 (12 days post-inoculation), macaque 31721 (56 days post-inoculation), and macaque 31816 (56 days post-inoculation) reads were mapped to each of the seven amplicons individually. 10,000 mapped reads were randomly subsampled from each amplicon in each sample, normalizing coverage at approximately 1,000x per nucleotide site throughout the complete genome. The normalized reads were then mapped to the consensus sequence of the inoculum using the bbmap read mapper (https://jgi.doe.gov). Minor variants that comprise at least 5% of total sequences in any of the four samples were identified using the bbmap callvariants.sh tool. The predicted effect of these variants on SWBV was calculated using snpEff (http://snpeff.sourceforge.net/) (Table S2). These sequencing datasets and Jupyter notebooks that create the data analysis environment and reproducibly perform the data analyses can be downloaded from https://go.wisc.edu/2k16q4.

### Viral loads

A TaqMan quantitative RT-PCR (RT-qPCR) assay was developed to quantify SWBV-1 plasma viral RNA (forward primer: 5’-GCTTGCTGGTAAGATTGCCA-3’; reverse primer: 5’-GCAGCGGATCTTTGTGGAA-3’; probe: 5’-6FAM-TGATTAACCTGAGGAAGTATGGCTGGC-BHQ1-3’), as described in detail previously (20). Briefly, RNA was extracted from 100 μl of plasma using the Viral Total Nucleic Acid Purification Kit (Promega, Madison, WI) on a Maxwell 16 MDx instrument and eluted in 50 μl. Reverse transcription and PCR were performed using the SuperScript III One-Step qRT-PCR system (Invitrogen, Carlsbad, CA) on a LightCycler 480 (Roche, Indianapolis, IN). Reverse transcription was carried out at 37°C for 15 minutes and then 50°C for 30 minutes followed by 2 minutes at 95°C, and then 50 cycles of amplification as follows: 95°C for 15 sec and 60°C for 1 minute. The 50 μl reaction mixture contained 5 μl of extracted RNA, MgSO_4_ at a final concentration of 3.0 mM, with the two amplification primers at a concentration of 500 nM and probe at a concentration of 100 nM. RNA copy numbers were calculated using a standard curve (described previously in (20)) that was sensitive down to 10 copies of RNA transcript per reaction.

### Peptide Array

Viral protein sequences were selected and submitted to Roche Sequencing Solutions (Madison, WI) for development into a peptide microarray as part of an early access program. Protein sequences were predicted from the SWBV inoculum nucleotide sequence and 16-mer peptides tiling each protein were designed to span each viral protein with a step size of 1 (amino acid overlap of 15). The peptide sequences were synthesized *in situ* with a Roche Sequencing Solutions Maskless Array Synthesizer (MAS) by light-directed solid-phase peptide synthesis using an amino-functionalized plastic support (Greiner Bio-One, Kremsmünster, Austria) coupled with a 6-aminohexanoic acid linker and amino acid derivatives carrying a photosensitive 2-(2-nitrophenyl) propyloxycarbonyl (NPPOC) protection group (Orgentis Chemicals, Gatersleben, Germany). To reduce instrument variance, each peptide was printed at 5 locations on the array. Samples were diluted 1:100 in binding buffer (0.01M Tris-Cl, pH 7.4, 1% alkali-soluble casein, 0.05% Tween-20). Diluted sample aliquots and binding buffer-only negative controls were bound to arrays overnight at 4°C. After binding, the arrays were washed 3× in wash buffer (1× TBS, 0.05% Tween-20), 10 min per wash. Primary sample binding was detected via 8F1-biotin mouse anti-primate IgG (NIH nonhuman primate reagent resource) secondary antibody. The secondary antibody was diluted 1:10,000 (final concentration 0.1 ng/μl) in secondary binding buffer (1× TBS, 1% alkali-soluble casein, 0.05% Tween-20) and incubated with arrays for 3 hours at room temperature, then washed 3× in wash buffer (10 min per wash) and 30 s in reagent-grade water. The secondary antibody was labeled with Cy5-Streptavidin (GE Healthcare; 5 ng/μl in 0.5x TBS, 1% alkali-soluble casein, 0.05% Tween-20) for 1 hour at room temperature, then the array was washed 2× for 1 min in 1× TBS, and washed once for 30 s in reagent-grade water. Fluorescent signal of the secondary antibody was detected by scanning at 635 nm at 2 μm resolution and 25% gain, using an MS200 microarray scanner (Roche NimbleGen, Madison, WI). The raw fluorescence signal intensity values were log2 transformed and the median transformed intensity of the redundant amino acid peptides was determined. Next, background subtraction was performed using the blank samples as background reactivity. Fold change from 0 DPI was calculated by subtracting the respective 0 DPI reactivity on a per amino acid peptide and per animal basis. The peptide array datasets and Jupyter notebooks that create the data analysis environment and reproducibly perform the data analyses can be downloaded from https://go.wisc.edu/83u95t.

### Flow cytometry

Cryopreserved peripheral blood mononuclear cells (PBMCs) were thawed at 37°C and washed in R10 media (RPMI containing 10% fetal bovine serum [FBS]). Between 3–5 million washed cells were transferred to 1.2 ml cluster tubes, and resuspended in phosphate-buffered saline (PBS) supplemented with 10% FBS (fluorescence-activated cell sorting [FACS] buffer). Cells were stained at 37°C with a mastermix of antibodies against CD38 (clone AT1, FITC conjugate, 20 μl), CD3 (clone SP34-2, Alexa Fluor 700 conjugate, 3 μl), CD8 (clone SK1, Brilliant Violet 510 conjugate, 2.5 μl), CD20 (clone 2H7, Brilliant Violet 650 conjugate, 2 μl), and CD4 (clone L200, Brilliant Violet 711 conjugate, 5 μl) antigens. All antibodies were obtained from BD BioSciences (San Jose, CA, USA), except the CD38-specific antibody, which was purchased from Stemcell Technologies (Vancouver, BC, Canada). Cells were also stained with LIVE/DEAD Fixable Near-IR during this time (Invitrogen, Carlsbad, CA). Cells were washed twice with FACS buffer, after which they were fixed with 0.125 mL of 2% paraformaldehyde for 20 min. After an additional wash the cells were permeabilized using Bulk Permeabilization Reagent from Invitrogen (Carlsbad, CA, USA). The cells were stained for 15 min with an antibody against Ki67 (clone B56, Alexa Fluor 647 conjugate, 5 μL) in the presence of permeabilizer. The cells were then washed twice and resuspended in 0.125 mL of 2% paraformaldehyde for an additional 20 min. After a final wash and resuspension with 0.125 mL FACS buffer, all samples were run on a BD LSRII Flow Cytometer within 24 hrs. Flow data were analyzed using Flowjo version 9.8.2.

### Statistical analysis

Information on statistical tests used to determine significance can be found in corresponding figure legends. All statistical analyses except those involving sequence analysis and peptide array analysis were performed using Prism7 (GraphPad Software, La Jolla, CA).

### Data accessibility

The sequence of the SWBV-1 inoculum used in this study can be found in GenBank (#MH686150). SWBV sequencing reads pertaining to this study can be found online at http://www.ncbi.nlm.nih.gov/bioproject/483240 under submission SRP155732 containing the accessions: SAMN09729057, SAMN09729058, SAMN09729059, and SAMN09729060. The workflow used to perform sequence analysis can be found at https://go.wisc.edu/2k16q4.

## Acknowledgements

This investigation was supported by a Pilot Grant from the Southwest National Primate Research Center grant P51OD011133 from the Office of Research Infrastructure Programs, National Institutes of Health. This work was also made possible by funding to the Wisconsin National Primate Research Center (P51RR000167) Office of Research Infrastructure Programs, National Institutes of Health. This research was conducted in part at a facility constructed with support from Research Facilities Improvement Program grant numbers RR15459-01 and RR020141-01. ALB performed this work with support from the University of Wisconsin– Madison’s Medical Scientist Training Program (MSTP) (grant T32GM008692) and the Physician Scientist Training Program (PSTP) sponsored by the Department of Pathology and Immunology at Washington University in St. Louis School of Medicine. The authors thank the University of Wisconsin, Department of Pathology and Laboratory Medicine and the WNPRC for funding and the use of its facilities and services. This work was also funded in part through the NIAID Division of Intramural Research (JPC and RFJ) and the Battelle Memorial Institute’s prime contract with the US National Institute of Allergy and Infectious Diseases (NIAID) under Contract No. HHSN272200700016I (JHK). This publication’s contents are solely the responsibility of the author(s) and do not necessarily represent the official views of ORIP, of ORIP, the US National Institutes of Health, NIAID, US Department of Health and Human Services, or of the institutions and companies affiliated with the authors. The funders of this research had no role in study design, data collection and analysis, decision to publish, or preparation of the manuscript. The authors have no conflicts of interest to declare. All authors edited and approved the manuscript.

